# Xeno-monitoring of molecular drivers of artemisinin and partner drug resistance in *P. falciparum* populations in malaria vectors across Cameroon

**DOI:** 10.1101/2022.01.19.476856

**Authors:** Francis N. Nkemngo, Leon M.J. Mugenzi, Magellan Tchouakui, Daniel Nguiffo-Nguete, Murielle J. Wondji, Bertrand Mbakam, Micareme Tchoupo, Cyrille Ndo, Samuel Wanji, Charles S. Wondji

## Abstract

**Background:** Monitoring of drug resistance in *Plasmodium* populations is crucial for malaria control. This has primarily been performed in humans and rarely in mosquitoes where parasites genetic recombination occurs. Here, we characterized the *Plasmodium* spp populations in wild *Anopheles* vectors by analyzing the genetic diversity of the *P. falciparum* kelch13 and *mdr1* gene fragments implicated in artemisinin and partner drug resistance across Cameroon in three major malaria vectors.

**Methods:** *Anopheles* mosquitoes were collected across nine localities in Cameroon and dissected into the head/thorax (H/T) and abdomen (Abd) after species identification. A TaqMan assay was performed to detect *Plasmodium* infection. Fragments of the Kelch 13 and *mdr1* genes were amplified in *P. falciparum* positive samples and directly sequenced to assess their drug resistance polymorphisms and genetic diversity profile.

**Results:** The study revealed a high *Plasmodium* infection rate in the major *Anopheles* vectors across Cameroon. Notably, A*n. funestus* vector recorded the highest sporozoite (8.02%) and oocyst (14.41%) infection rates. A high *P. falciparum* sporozoite rate (80.08%) alongside epidemiological signatures of significant *P. malariae* (15.94%) circulation were recorded in these vectors. Low genetic diversity with six (A578S, R575I, G450R, L663L, G453D, N458D) and eight (H53H, V62L, V77E, N86Y, G102G, L132I, H143H, Y184F) point mutations were observed in the *k13* and *mdr1* backbones respectively. Remarkably, the R575I (4.44%) *k13* and Y184F (64.2%) *mdr1* mutations were the predominant variants in the *P. falciparum* populations.

**Conclusion:** The emerging signal of the R575I polymorphism in the *Pfk13* propeller backbone entails the regular surveillance of molecular markers to inform evidence-based policy decisions. Moreover, the high frequency of the ^86^N^184^F allele highlights concerns on the plausible decline in efficacy of artemisinin-combination therapies (ACTs); further implying that parasite genotyping from mosquitoes can provide a more relevant scale for quantifying resistance epidemiology in the field.

## 1. Introduction

The triad biological interaction between *Anopheles* vectors, *Plasmodium* parasites, and humans is responsible for malaria, a disease of global public health priority [1]. The disease burden predominates in sub-Saharan Africa with Cameroon accounting for 6,450,000 cases in 2020, mostly affecting children below five years [1–3]. This high score in malaria cases is in part due to the heterogenous vectorial complexity and competence of the major *Anopheles* mosquitoes particularly *An. gambiae*, *An. funestus* and *An. coluzzii* in successfully transmitting the deadliest, most prevalent, most drug-resistant *Plasmodium* species, *P. falciparum* across diverse bio- ecological zones within the country [3–5]. *P. falciparum* is responsible for about 93% of cases and death with the remaining fraction attributed to non-falciparum species including *P. malariae, P. ovale,* and *P. vivax* (hereafter referred to as P. OVM) [6–8]. In particular, *P. malariae* significantly predominates as co-infection with *P. falciparum* to widen the parasite transmission window and disease severity [9]. *Anopheles* mosquito serves both as the vector and the definitive host of the *Plasmodium* parasite and this obligatory co-interaction forms the basis of sporogony [10, 11]. This stage, usually considered the most important in the parasite life cycle owing to the occurrence of fertilization, genetic exchange, allelic recombination, and sporozoite formation defines the cornerstone of malaria transmission [12, 13]. Gametocytes, which comprise 1% (∼100-1000) of the parasite biomass [14] are the sexual stages involved in the transmission of both drug-resistant and sensitive alleles. In line with this, antimalarial drugs have played a key role in shrinking the parasite population in humans while also preventing mosquitoes from being infected during a blood meal [15, 16]. In particular, Artemisinin combination therapy (ACT), the first-line treatment against *P. falciparum* infection, has contributed enormously both in decreasing the parasite biomass in humans and reducing transmission capacity [17, 18]. Indeed, a modeling study between 2000 and 2015 revealed that 19% of the success in averting malaria burden in endemic countries was attributed to expanded access to ACTs [19]; diminishing the global malaria mortality curve from 896,000 deaths in 2000 to 627,000 deaths in 2020 [1]. However, the emergence of the artemisinin-resistant (AR) *P. falciparum* parasite in Western Cambodia (South East Asia) and recently in Rwanda and Uganda (Africa) threatens to reverse the gains achieved in malaria control over the years [20–23]. Moreover, the widespread molecular resistance to the partner drugs (amodiaquine and mefloquine) is jeopardizing malaria control efforts in the most affected regions [1]. This poses an even greater risk to the clinical therapeutic efficacy and sustainability of ACTs particularly in Africa in the absence of newly approved compounds [24]. In order to mitigate the potential risks of Artemisinin and partner drug resistance, molecular markers have been identified to detect resistance at the early stage to improve treatment schemes.

In this regard, the discovery of molecular markers of resistance in the Kelch 13 (*k13*) [25] and multidrug resistance-1 (*mdr1*) [26] genes through whole-genome sequencing approach and phenotypic studies has facilitated the possibility to monitor and track the emergence and spread of artemisinin and partner drug resistance in real-time and geography. *Plasmodium falciparum* drug resistance monitoring is critical for successful malaria control and elimination [1]. Techniques involving genotyping of resistance markers from human blood samples are usually invasive, require huge logistic costs, are time-consuming, and involve a lengthy duration of ethical considerations particularly in malaria-endemic countries [22]. An alternative cost-effective approach to surveillance of resistant parasites from human blood in endemic areas is utilizing *Plasmodium*-infected *Anopheles* mosquitoes as a sentinel for the screening of drug resistance genes [27].

Exploiting *Plasmodium*-infected mosquitoes to detect and track resistant parasites in endemic areas through genotyping of molecular markers or assessing signatures of selection through reduced diversity, could provide an early warning signal for the emergence of drug resistance [27, 28]. In addition, it will permit the rapid follow-up of the geographical distribution and spread of drug resistance within a country thereby triggering pro-active responses to improve local malaria control strategies [29].

Therefore, this study aims to firstly establish the prevalence of *Plasmodium* infection in *Anopheles* vectors across nine (09) localities in Cameroon and secondly to characterize the polymorphism profile and genetic variability of *P. falciparum k13* and *mdr1* gene determinants implicated in artemisinin and partner drug resistance in the dominant *Anopheles* mosquitoes circulating across Cameroon.

## 2. Methodology

### 2.1. Study sites

This study was conducted across 09 localities in Cameroon (Fig. 1) representing three major bio- ecological belts mainly equatorial, sudano-guinean, and Sahel regions. The study sites categorized within the tropical equatorial region includes Bankeng (4° 38 43 N; 12° 13 03 E) ″ ″ [30], Bonaberi (4° 4’ 55.955”N; 9° 39’ 53.898” E) [5], Elende (3°41’57.27’’N, 11°33’28.46’’E) [2], Elon (N4.23051° E11.60120°) [31], Obout (3’ 7’0 ”N, 11’ 65’0” E) [32] and Mangoum (5°31’N, 10°37’E) [5]. These rural areas (except Bonaberi, an urbanized locality) situated within the forested parts of Cameroon have a high humidity profile characterized by two rainy season shifts, ranging from March to June and September to November. Malaria transmission in this climatic zone is considered stable and perennial. Moreover, the vector dominance of *An. gambiae* s.s in Bankeng and Mangoum; *An. coluzzii* in Bonaberi and *An. funestus* in Elende, Elon, and Obout pilots the malaria transmission pattern observed in these areas [2, 5, 30, 31]. 59 E) situated in the sudano- ″ guinean Adamawa region, forms a mid-point between the tropical equatorial south and the savanna north. The climate is characterized by a rainy season from May to September, and a dry season extending from October to April. *An. funestus* and *An. coluzzii* dominate in Mibellon and Gounougou respectively [5, 33, 34]. Simatou (10°34′ ′ characterized by a short periodic rainy season from May to September and a long dry season from October to April [5]. Malaria transmission in this area is seasonal with *An*. *coluzzii* being the leading vector.

**Fig. 1.**
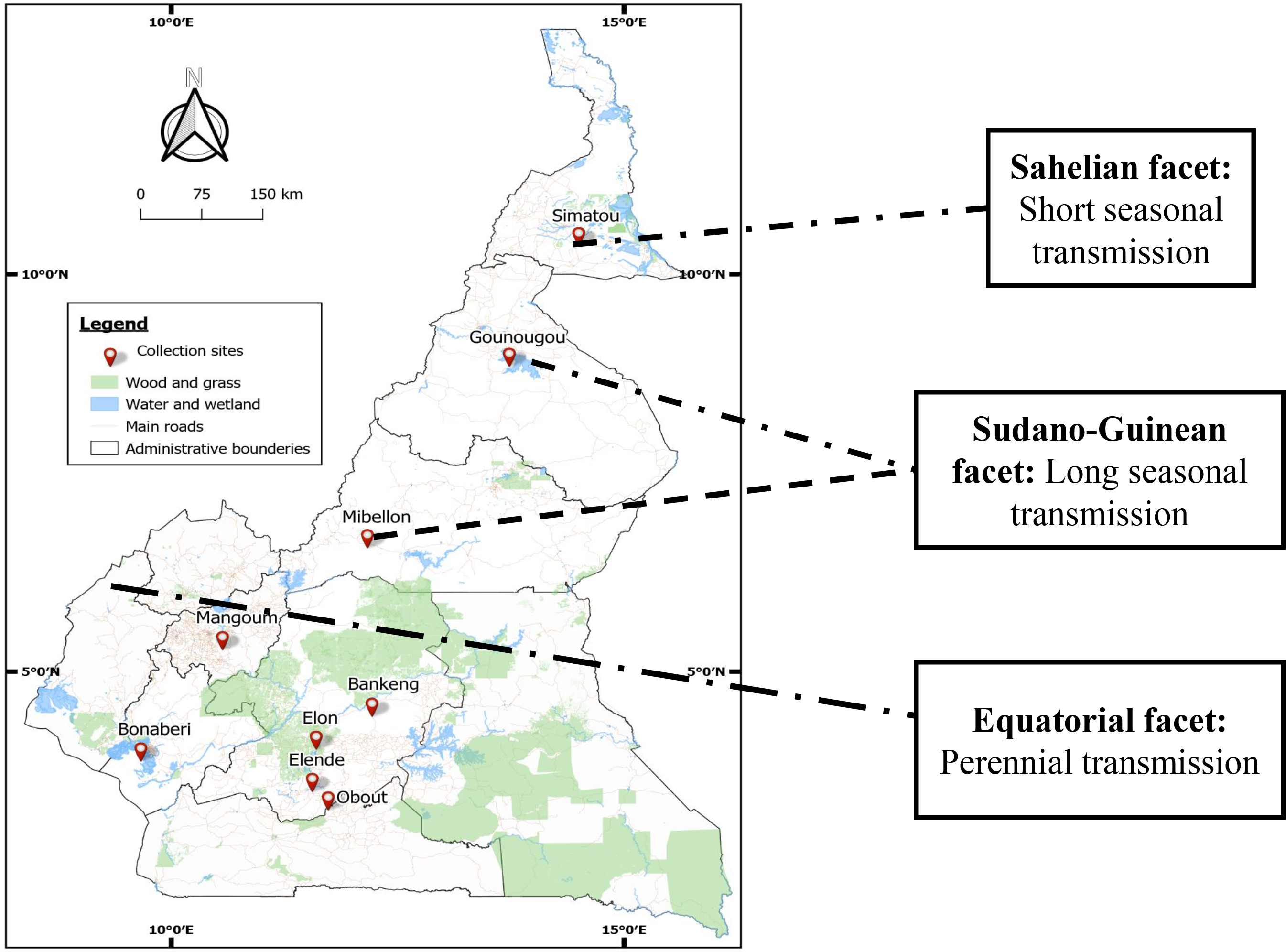

### 2.2. Study design: Mosquito collection across Cameroon

Adult mosquitoes were collected across the 09 localities (Fig. 1) from houses using different collection techniques after ethical approval was obtained from the Cameroon National Committee on Research Ethics for Human Health (N^0^ 2020/05/1234/CE/CNERSH/SP) and verbal consent was sought from village heads and household members. Indoor resting mosquito populations from Elende, Elon, Mibellon, Obout were collected on the walls and roofs of thatched houses using electrical aspirators (Rule In-Line Blowers, Model 240) [2] while Human Landing Catch (HLC) was employed to capture mosquitoes both inside and outside of homes in the localities of Bonaberi and Gounougou [5]. In Bankeng, both indoor aspiration and HLC techniques were utilized for mosquito collection while HLC and Pyrethrum Spray Catch (PSC) methods were used for mosquito sampling in Mangoum and Simatou [5]. The period of mosquito collection varied across the different study sites: Bankeng (April 2019), Bonaberi (April, August & October 2019), Elende (April to June 2019), Elon (April 2019 & January 2020), Gounougou (April 2019 & June 2020), Mangoum (April 2019 & October 2020), Mibellon (August to

September 2019), Obout (May 2016) and Simatou (April 2019, March & June-July 2020). All mosquitoes (F0) were morphologically identified as either *An. gambiae* complex or *An. funestus* group following established protocols [35, 36].

Female *Anopheles* mosquito abdomen (Abd) and head/thorax (H/T) were partitioned to discriminate between midgut (Abdomen) and salivary gland (H/T) infection [37]. However, this separation technique was not done for mosquito samples from Obout as collected back in 2016. Rather, whole mosquitoes (WM) were used for extraction. All the samples were stored at -20°C until DNA extraction.

Extraction of genomic DNA from the H/T and Abd of individual dissected female mosquitoes from all the localities (except Obout) was accomplished using the LIVAK method [38]. Species identification by PCR was performed to discriminate against *An. funestus* siblings [39] while the SINE-200 method was employed to distinguish members of the *An. gambiae* s.l species complex [40].

### 2.3. TaqMan detection of *Plasmodium* sporozoites and oocysts in field-collected *Anopheles* mosquitoes

Screening for *Plasmodium* infection was done to check for the presence of sporozoite and oocyst using the TaqMan assay [41] for nine localities. The number of samples tested per locality include: Bankeng (n= 287H/T), Bonaberi (n= 262H/T & 372Abd), Elende (n= 1000H/T & 434Abd), Elon (n= 273H/T & 378Abd), Gounougou (n= 465H/T & 558Abd), Mangoum (n= 465Abd), Mibellon (n= 640H/T & 372Abd), Obout (n= 186WM) and Simatou (n= 372H/T and 465Abd). The real-time PCR MX 3005 (Agilent, Santa Clara, CA, USA) system was used for the amplification [41]. Briefly, 2 μl of gDNA for each sample was used as template in a 3-step program with a pre-denaturation at 95°C for 10 mins, followed by 40 cycles of 15 sec at 95°C and 1 min at 60°C. The primers (Falcip+: TCT-GAA-TAC-GAA-TGT-C, OVM+: CTG-AAT- ACA-AAT-GCC, Plas-F: GCT-TAG-TTA-CGA-TTA-ATA-GGA-GTAGCT-TG, Plas R: GAA-AAT-CTA-AGA-ATT-TCA-CCTCTG-ACA) were used together with two probes tagged with fluorophores: FAM to detect *Plasmodium falciparum*, and HEX to detect *Plasmodium ovale*, *Plasmodium vivax,* and *Plasmodium malariae. P. falciparum* samples and a mix of *P. ovale*, *P. vivax,* and *P. malariae* were used as positive controls. A sub-set of positive samples (for each body part and locality) were subjected to Nested PCR to confirm and discriminate the species detected by TaqMan based on the protocol of [42] with slight modification using kappa Taq enzyme instead of Dream Taq.

### 2.4. Amplification of *k13* propeller domain and *mdr1* gene fragments of *P. falciparum* from infected *Anopheles* mosquitoes

The *P. falciparum-positive* DNA samples were used as templates to nested amplify a portion of the *kelch13* gene encompassing the propeller domain known to contain the key mutations mediating artemisinin resistance [25, 43]. The primary and nested PCR *k13* primers are found in Table 1: S1. For the primary PCR, 20 µL of the final volume was constituted of 4 µL of the genomic DNA extract; 0.51µL each forward and reverse primers; 0.12 µL each kappa Taq enzyme (Kappa Biosystems, Wilmington, MA, USA) and dNTP mix; 0.75 µL MgCl_2_, 1.5 µL kappa Taq buffer, and 12.49 µL distilled water. The thermocycling conditions include: initial denaturation at 95 ^0^C for 5 min, followed by 30 cycles each of 30 sec at 95 ^0^C (denaturation), 2 min at 58 ^0^C (primer annealing), 2 min sec at 72 ^0^C (elongation). This was followed by a 10 min final elongation at 72 ^0^C. The nested PCR followed the same protocol of master mix composition as the primary PCR. The thermocycling parameters were the same except for the annealing temperature at 59 ^0^C for 30 sec and 35 cycles increment.

**Table 1.** Status of Plasmodium infection rate in the major Anopheles malaria vectors across Cameroon: (**a**) Head/thorax (HT), (**b**) Abdomen (Abd), **(c)** Whole mosquito (WM); where: N= number of mosquito samples examined; Falcip+= infection by P. falciparum; OVM+= infection by P. ovale/vivax/malariae; and Falcip+/OVM+= Co- infection by Plasmodium falciparum and P. ovale/vivax/malariae; AGAM = An. gambiae s.s; AFUN = An. funestus s.s and ACOL = An. coluzzii.

| Localities | *Anopheles* spp | Year(s) of collection | N | Total infection | Falcip+ | OVM+ | Falcip+/OVM+ |
| --- | --- | --- | --- | --- | --- | --- | --- |
| **Head/Thorax: *Plasmodium* sporozoite infection rate** |  |  |  |  |  |  |  |
| Bankeng | *An. gambiae* | 2019 | 287 | 22 (7.67%)<br>[5.6-10.27%] | 20 (90.90%)<br>[87.37-94.71%] | 01 (4.55%)<br>[2.22-7.57%] | 01 (4.54%)<br>[2.22-7.57%] |
| Bonaberi | *An. coluzzii* | 2019 | 262 | 01 (0.38%)<br>[0.13-0.58%] | 01 (100%) | 00 (0%) | 00 (0%) |
| Elende | *An. funestus* | 2019 | 1000 | 78 (7.80%)<br>[6.80-11.11%] | 68 (87.17%)<br>[84.96-90.41%] | 08 (10.25%)<br>[8.06-13.46%] | 02 (2.56%)<br>[0.93-5.33%] |
| Elon | *An. funestus* | 2019 & 2020 | 273 | 07 (2.56%)<br>[1.36-4.54%] | 07 (100%) | 00 (0%) | 00 (0%) |
| Gounougou | *An. coluzzii* | 2019 & 2020 | 465 | 34 (7.31%)<br>[6.58-12.03%] | 23 (67.65%)<br>[65.26-70.76%] | 09 (26.47%)<br>[23.48-28.55%] | 02 (5.88%)<br>[3.27-8.64%] |
| Mibellon | *An. funestus* | 2019 | 640 | 88 (13.75%)<br>[12.11-20.11%] | 62 (70.45%)<br>[67.13-73.44%] | 21 (23.86%)<br>[20.58-25.03%] | 05 (5.68)<br>[3.39-7.06%] |
| Simatou | *An. coluzzii* | 2019 & 2020 | 372 | 21 (5.65)<br>[4.20-8.45%] | 20 (95.24%)<br>[92.65-97.94%] | 01 (4.76%)<br>[2.14-6.42%] | 00 (0%) |
| **Abdomen: *Plasmodium* oocyst infection rate** |  |  |  |  |  |  |  |
| Bonaberi | *An. coluzzii* | 2019 | 372 | 04 (1.08%)<br>[0.67-2.67%] | 04 (100%) | 00 (0%) | 00 (0%) |
| Elende | *An. funestus* | 2019 | 434 | 47 (10.83%)<br>[8.93-15.37%] | 44 (93.62%)<br>[90.52-95.48%] | 03 (6.38%)<br>[3.68-9.42%] | 00 (0%) |
| Elon | *An. funestus* | 2019 | 378 | 28 (7.41%)<br>[7.25-9.04%] | 26 (92.86%)<br>[89.03-95.96%] | 02 (7.14%)<br>[5.43-9.63%] | 00 (0%) |
| Gounougou | *An. coluzzii* | 2019 & 2020 | 558 | 29 (5.20%)<br>[3.83-8.83%] | 20 (68.97%)<br>[65.98-70.42%] | 07 (24.14%)<br>[21.87-27.69%] | 02 (6.89%)<br>[4.15-8.54%] |
| Mangoum | *An. gambiae* | 2019 & 2020 | 465 | 51 (10.97%)<br>[9.93-12.99%] | 46 (90.20%)<br>[87.96-92.03%] | 05 (9.80%)<br>[7.02-11.29%] | 00 (0%) |
| Mibellon | *An. funestus* | 2019 | 372 | 93 (25.00%)<br>[23-27%] | 56 (60.22%)<br>[58.24-62.07%] | 35 (37.63%)<br>[35.87-40.45%] | 02 (2.15%)<br>[0.92-4.54%] |
| Simatou | *An. coluzzii* | 2019 & 2020 | 465 | 25 (5.38%)<br>[5.22-5.55%] | 21 (84%)<br>[81-86%] | 04 (16%)<br>[14-18%] | 00 (0%) |
| **Whole mosquito: *Plasmodium* infection rate** |  |  |  |  |  |  |  |
| Obout | *An. funestus* | 2016 | 186 | 72 (38.71%)<br>[35.7-41.36%] | 57 (79.17%)<br>[76.1-82.56%] | 09 (12.5%)<br>[10.03-14.48%] | 06 (8.33%)<br>[6.22-10.44%] |

Similarly, a nested PCR approach was used for amplification of codons 86 and 184 fragments of the *mdr1* gene [44]. The primary and nested *mdr1* PCR primers are included in Table 1: S1. The traditional primary and nested *mdr1* PCR master mix composition was similar to that of the *k13* protocol. The thermocycling profile was: initial denaturation at 95 ^0^C for 5 min, proceeded by 30 cycles each of 30 sec at 94 ^0^C, 45 sec at 45 ^0^C, 1 min at 72 ^0^C, and a final elongation of 10 min for 72 ^0^C. The nested PCR protocol was the same like the primary except for the annealing temperature at 52 ^0^C for 45 sec and 35 cycles addition. After amplification, the nested PCR products of both genes were each separated in 2% agarose gel. Furthermore, 10 µL PCR products each of the samples that correctly amplified were purified by the Exo-SAP protocol [New England Biolabs (NEB, MA, and USA)]. At most 20 (range: 4-20) randomly selected amplified *Pfk13* and *Pfmdr1* samples each from the HT and Abd body parts (depending on the infection rate) of the *Anopheles* mosquitoes from all the localities were directly sequenced.

### 2.5. Data analysis

The sporozoite and oocyst infection rates for the major *Anopheles* vectors across the 09 localities in Cameroon were analyzed in GraphPad Prism V8 (GraphPad Software, La Jolla California USA). A Chi-square test was used to determine the differences between categorical variables of stage-specific *Plasmodium* infection prevalence in each *Anopheles* vector per the locality. On the other hand, genomic sequence analysis commenced with a visual inspection of the qualities of the DNA sequence chromatograms and FASTA file using Chromas V.2.5 and Bioedit V.7.2.5 software respectively [45]. The trimmed sequences were aligned to the *Pfk13* (PF3D7_1343700) and *Pfmdr1* (PF3D7_0523000) reference sequences in PlasmoDB (www.Plasmodb.org) and examined for polymorphisms using ClustalW tool. A consensus forward sequence for each parasite population according to the body part (HT, Abd and Whole) and *Anopheles* species was generated with Bioedit software. Nucleotide sequences were translated *in silico* to complementary amino acids using the appropriate open reading frame in Mega X version 10.1.6 [46] to identify the relevant single nucleotide polymorphism (SNP). A cladogram was built using the Maximum Likelihood method and Tamura-3 model, with a bootstrap factor of 1000 replicates. DNA polymorphisms were generated in dnaSP V.6.12.03 [47]. Haplotype networks were constructed using a combination of Arlequin 3.5.2.2. [48] and PopART [49] software.

## 3. Results

### 3.1. Species identification

Extraction of genomic DNA was done for a total of 6529 individual *Anopheles* mosquito samples (HT, Abd, and WM) across 09 localities in Cameroon (Table 1). Molecular speciation on a sub- set of 93 randomly extracted mosquito DNA samples across the 09 localities confirmed the dominance of *An. gambiae* s.s. in Bankeng [98.9% (92/93)] and Mangoum [100% (93/93)], *An. coluzzii* in Bonaberi [97.9% (91/93)], Gounougou [81.7% (76/93)] and Simatou [86.0% (80/93)]; and *An. funestus* s.s in Elende [97.8% (91/93)], Elon [93.5% (87/93)], Mibellon [98.9% (92/93)] and Obout [100% (93/93)].

### 3.2. Comparative *Plasmodium* infection rate in the head/thorax, abdomen, and whole mosquito of the major malaria vectors

The analysis of the head and thorax (HT) and abdomen (Abd) of the *Anopheles* mosquitoes across the 09 localities reveals a varying *Plasmodium* sporozoite and oocyst infection rate (Table 1; Fig. 2). Generally, sporozoite infection rates ranged from 0.38% (1/262) in *An. coluzzii* to 7.67% in *An. gambiae* and 13.75% (88/640) in *An. funestus.* The predominant species was *P. falciparum* with a frequency ranging from 67.65% to 100%%. Meanwhile, OVM+ vary from 0% to 26.47% and mix infection spanned from 0% to 5.88% (Table 2). Similar pattern was observed for oocyst infection rates although at a higher level. This ranges from 1.08% (4/372) in *An. coluzzii* to 25% (93/372) in *An. funestus. P. falciparum* was the main species with a frequency ranging from 60.22% to 100%. Likewise, the OVM+ infection ranges from 0% to 37.63% while mix infection vary from 0% to 6.89% (Table 1). In whole body *An. funestus* samples from Obout, the *Plasmodium* infection rate was 38.71% (72/186) with 79.17% *P. falciparum*, 12.5% OVM+ and 8.33% mix infection.

**Fig. 2.**
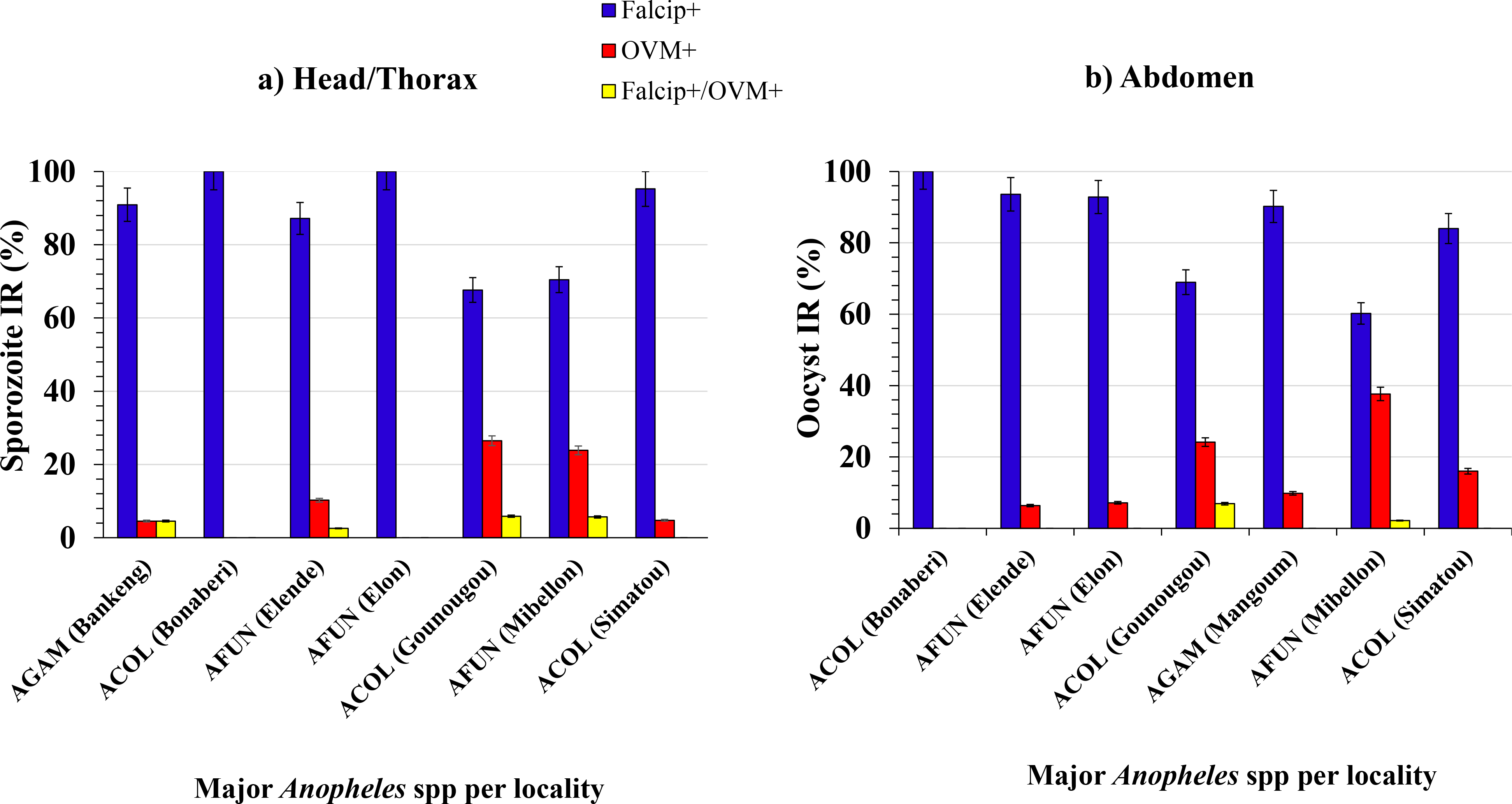

**Table 2.** Key synonymous (S) and non-synonymous (NS) single-nucleotide polymorphisms in P. falciparum drug resistance markers from infected Anopheles mosquitoes across Cameroon: **a.** k13 β propeller domain and **b.** mdr1gene fragments **Note:** The boldface highlights the nucleotide base change. **Abbreviations:** n = number of samples containing mutant allele; N = total number of successfully sequenced samples; NS = non-synonymous mutation; S = synonymous mutation; ***Pfk13* polymorphisms**: A = Alanine, S = Serine, G = Glycine, R = Arginine, L= Leucine, D = Aspartic acid, I = Isoleucine, N = Asparagine. ***Pfmdr1* polymorphisms**: Y = Tyrosine, F = Phenylalanine, V = Valine, H = Histidine

| Gene | Body part | *Anopheles* spp | Codon position | Wild sequence |  | Mutant sequence |  | Amino acid Product | Type | Frequency (%) | n/N |
| --- | --- | --- | --- | --- | --- | --- | --- | --- | --- | --- | --- |
|  |  |  |  | Amino acid | Nucleotide | Amino acid | Nucleotide |  |  |  |  |
| *Pfk13* | Abdomen (Midgut) | AGAM | 1348 | G | GGA | R | AGA | G450R | NS | 2.22 | 1/45 |
|  |  | AFUN | 1358 | G | GGT | D | GAT | G453D | NS | 2.22 | 1/45 |
|  |  |  | 1724 | R | AGA | I | ATA | R575I | NS | 4.44 | 2/45 |
|  |  |  | 1732 | A | GCT | S | TCT | A578S | NS | 2.22 | 1/45 |
|  |  |  | 1987 | L | CTA | L | TTA | L663L | S | 4.44 | 2/45 |
|  | HT (Salivary gland) | AFUN | 1372 | N | AAT | D | GAT | N458D | NS | 2.94 | 1/34 |
|  |  |  | 1732 | A | GCT | S | TCT | A578S | NS | 2.94 | 1/34 |
|  |  |  | 1987 | L | CTA | L | TTA | L663L | S | 2.94 | 1/34 |

**Table 2.** Key synonymous (S) and non-synonymous (NS) single-nucleotide polymorphisms in *P. falciparum* drug resistance markers from infected *Anopheles* mosquitoes across Cameroon: **a.** *k13* $\beta$ propeller domain and **b.** *mdr1* gene fragments
| *Pfmdr1* |  |  |  |  |  |  |  |  |  |  |
| --- | --- | --- | --- | --- | --- | --- | --- | --- | --- | --- |
| *Anopheles* vector | N | Oocyst stage (Abdomen) |  |  |  |  |  |  |  |  |
|  |  | V62L<br>n (%) |  |  | N86Y<br>n (%) |  |  | Y184F<br>n (%) |  |  |
|  |  | Wild<br>(62V) | Mutant<br>(62L) | Mixed<br>(62V/L) | Wild<br>(86N) | Mutant<br>(86Y) | Mixed<br>(86N/Y) | Wild<br>(184Y) | Mutant<br>(184F) | Mixed<br>(184Y/F) |
| ACOL | 29 | 28 (96.6) | 1(3.5) | 0 (0) | 26 (89.7) | 1 (3.5) | 2 (6.9) | 10 (34.5) | 15 (51.7) | 4 (13.8) |
| AGAM | 16 | 16 (100) | 0 (0) | 0 (0) | 13 (81.3) | 1 (6.3) | 2 (12.5) | 0 (0) | 8 (50) | 8 (50) |
| AFUN | 48 | 42 (87.5) | 5 (10.4) | 1 (2.1) | 40 (83.3) | 3 (6.3) | 5 (10.4) | 9 (18.8) | 33 (68.8) | 6 (12.5) |
| All | 93 | 86 (92.5) | 6 (6.4) | 1 (1.1) | 79 (84.9) | 5 (5.4) | 9 (9.7) | 19 (20.4) | 56 (60.2) | 18 (19.4) |
| **Sporozoite stage (H/T)** |  |  |  |  |  |  |  |  |  |  |
| ACOL | 23 | 22 (95.7) | 1 (4.3) | 0 (0) | 15 (65.2) | 4 (18.2) | 4 (17.4) | 3 (13.0) | 19 (82.6) | 1 (4.3) |
| AGAM | 17 | 17 (100) | 0 (0) | 0 (0) | 11 (64.7) | 3 (17.6) | 3 (17.6) | 3 (17.6) | 11 (64.7) | 3 (17.6) |
| AFUN | 40 | 39 (97.5) | 1 (2.5) | 0 (0) | 29 (72.5) | 7 (17.5) | 4 (10) | 7 (17.5) | 24 (60) | 9 (22.5) |
| All | 80 | 78 (97.5) | 2 (2.5) | 0 (0) | 55 (68.8) | 14 (17.5) | 11 (13.8) | 13 (16.3) | 54 (67.5) | 13 (16.3) |
| **Total Plasmodium (Whole body)** |  |  |  |  |  |  |  |  |  |  |
| AFUN | 20 | 0 (0) | 0 (0) | 6 (30) | 13 (65) | 1 (5) | 6 (30) | 3 (15) | 13 (65) | 4 (20) |

Overall, *An. funestus* population from Mibellon exhibited the highest *P. falciparum* and *P. malariae* sporozoite and oocyst infection rates while the least infection rate was recorded in *An. coluzzii* population from Bonaberi (Table 1). Moreover, a significant difference was observed between mosquitoes infected with *Plasmodium* oocyst and those with the infective sporozoite stage in Mibellon (2 = 2.44; P LJ 0.05), Elende (2 = 3.85; P LJ 0.05) and Elon (2 = 5.62; P LJ 0.05). In addition, mosquitoes collected using the techniques of indoor aspiration (*Plasmodium* Infection Rate (PIR) = 18.78%) and pyrethrum spray catch (PSC) (PIR = 7.09%) yielded a high *Plasmodium* infection rate than mosquitoes collected by HLC (PIR = 3.87%) (Table 2: S1).

### 3.3. Detection of polymorphisms in the *Pf*kelch13 propeller domain (*k13*PD) and *mdr1* gene fragments

An 830bp fragment of *Pfk13* gene encompassing the β 789 coding sequence) was successfully amplified (Fig 1a: S1) from 171 *Pfk13* samples (88 Abd, 69 H/T and 14 WM) (Table 3a: S1). Out of the 171 *P. falciparum* sequences, 158 were similar to the 3D7 reference. Three unique point mutations were observed in four oocyst-infected *Anopheles* abdomen sequences including G450R (2.22%) in *An. gambiae*, G453D (2.22%) and L663L (4.44%) in *An. funestus* meanwhile the N458D (2.94%) and L663L (2.94%) were observed in two sporozoite infected H/T *An. funestus* sequences (Table 2a). The A578S mutation was present in both the oocyst and sporozoite sequences of *An. funestus*. Also, the R575I (4.44%) was present in two *P. falciparum* oocyst sequences of *An. funestus* but absent at the sporozoite stage (Table 2a). In addition, ambiguous polymorphisms were present only in *P. falciparum* oocyst sequences including L462(L/L) in *An. gambiae* and S466(S/T) in *An. funestus*. None of the WHO-validated or candidate *Pfk13* polymorphisms associated with artemisinin resistance were identified (Table 2a).

**Table 3a.** Polymorphism and genetic diversity parameters of k13 drug resistance marker in natural P. falciparum populations circulating in the head/thorax and abdomen of major Anopheles vectors across Cameroon. **Abbreviations:** n = number of sequences; S = number of polymorphic sites; H = haplotype; Hd = haplotype diversity; = nucleotide diversity; TajimaD = Tajima’s D statistic; FuLiD* = Fu and Li’s D* statistic; FuLiF* = Fu and Li’s F* statistic; * = significant. AGAM = An. gambiae s.s; AFUN = An. funestus s.s and ACOL = An. coluzzii. 2n* = Sequences were unphased because of the observed heterozygosity (note: P. falciparum is haploid during the sporozoite stage (HT) of the mosquito. However, manual observation of the H/T mdr1 sequences revealed a high heterozygosity. Thus, the sequences were unphased to have a detailed picture of the mixed alleles present in the sequences. This observed heterozygosity could be due to parasites originating from different tribal lineages or as a result of the high copy number variation of this gene).

| Body part | Species | Gene | 2(n) | S | H | Hd | Pi | TajimaD | FuLiD | FuLiF |
| --- | --- | --- | --- | --- | --- | --- | --- | --- | --- | --- |
| Abdomen<br>(Diploid oocyst); 2n | ACOL | *k13* | 56 | 0 | 1 | 0 | 0 | 0 | 0 | 0 |
|  | AGAM |  | 30 | 2 | 3 | 0.191 | 0.00025 | -1.25553 | -0.73747 | -1.02054 |
|  | AFUN |  | 90 | 6 | 8 | 0.248 | 0.00041 | -1.65933 | -0.75796 | -1.24546 |
|  | All |  | 176 | 8 | 10 | 0.163 | 0.00026 | -1.91573* | -1.39175 | -1.87876 |
| Head/Thorax<br>(Haploid sporozoite); 2n* | ACOL |  | 40 | 1 | 2 | 0.050 | 0.00006 | -1.12411 | -1.77404 | -1.83507 |
|  | AGAM |  | 30 | 0 | 1 | 0 | 0 | 0 | 0 | 0 |
|  | AFUN |  | 68 | 4 | 5 | 0.195 | 0.00026 | -1.61083 | -0.17526 | -0.73353 |
|  | All |  | 138 | 5 | 6 | 0.113 | 0.00015 | -1.76645 | -1.19624 | -1.63960 |
| Abdomen (oocyst)<br>+ Head/Thorax<br>(sporozoite) | ACOL |  | 96 | 1 | 2 | 0.021 | 0.00003 | -1.03241 | -2.02060 | -2.00827 |
|  | AGAM |  | 60 | 2 | 3 | 0.098 | 0.00013 | -1.31528 | -0.94290 | -1.22624 |
|  | AFUN |  | 158 | 8 | 10 | 0.225 | 0.00035 | -1.83601* | -1.34976 | -1.80964 |
|  | All |  | 314 | 11 | 13 | 0.141 | 0.00021 | -2.05516* | -2.52346* | -2.82689* |
| Whole mosquito;<br>2n* (mixed oocyst<br>+ sporozoite stages) | AFUN |  | 28 | 0 | 1 | 0 | 0 | 0 | 0 | 0 |

Similarly, a 570-base pair of the *mdr1* gene portion covering locus 86 to 184 (ranging from nucleotide position 121 to 597) was amplified from DNA extracts of *P. falciparum* infected HT, Abd and whole mosquito samples across the various localities (Fig 1b: S1). A total of 193 *Pfmdr1* samples (93 Abd, 80 HT and 20 WM) were sequenced successfully (Table 3b: S1). Generally, the prevalence of circulating *P. falciparum* parasites harboring the mutant N86Y and Y184F non-synonymous polymorphisms were 5.4% and 60.2% in the oocyst stage; 17.5% and 67.5% in the sporozoite and 5% and 65% in the mixed stages from whole body respectively (Table 2b). Contrasting frequencies of the 86Y and 184F alleles were observed (Fig. 3 & Table 2b), indicative of the selection created by the major ACT, notably Artemether-Lumefantrine (AL) on the parasite genome. The presence of the double mutant haplotype YF was 5.4%, 16.3% and 5% in the oocyst, sporozoite and mixed stages from whole body respectively, with no significant difference observed across the different *Anopheles* populations. Additionally, the single mutant haplotype NF was the most predominant with a prevalence of 45.2%, 36.3% and 30% in the oocyst, sporozoite and mixed stages from whole body respectively. The NY haplotype occurred at a frequency of 20.4%, 16.3% and 15% in the oocyst, sporozoite and mixed stages from whole body respectively while the YY haplotype was absent (Table 4b: S1). Interestingly, a novel emerging variant, V62L was observed at low frequency in the abdomen (6.4%) and H/T (2.5%) alongside

**Table 3b.** Polymorphism and genetic diversity parameters of *mdr1* codons 86 & 184 drug resistance marker in natural *P. falciparum* populations circulating in the head/thorax, abdomen and whole body of the major *Anopheles* vectors across Cameroon. **Abbreviations:** n = number of sequences; S = number of polymorphic sites; H = haplotype; Hd = haplotype diversity; π = nucleotide diversity; TajimaD = Tajima’s D statistic; FuLiD* = Fu and Li’s D* statistic; FuLiF* = Fu and Li’s F* statistic; * = significant; 2n* = Sequences were unphased because of the observed high heterozygosity associated with gene copy number variation. AGAM = *An. gambiae* s.s; AFUN = *An. funestus* s.s and ACOL = *An. Coluzzii*

| Body part | Species | Gene | 2(n) | S | H | Hd | Pi | TajimaD | FuLiD | FuLiF |
| --- | --- | --- | --- | --- | --- | --- | --- | --- | --- | --- |
| Abdomen<br>(Diploid oocyst); 2n | ACOL | mdr1 | 58 | 5 | 7 | 0.656 | 0.00187 | -0.77269 | -0.60065 | -0.76994 |
|  | AGAM |  | 32 | 3 | 4 | 0.573 | 0.00154 | -0.03430 | 0.94181 | 0.76400 |
|  | AFUN |  | 96 | 4 | 6 | 0.618 | 0.00186 | -0.19123 | 1.04021 | 0.75155 |
|  | All |  | 186 | 6 | 10 | 0.626 | 0.00183 | -0.59191 | 1.14595 | 0.64412 |
| Head/Thorax<br>(Haploid sporozoite); 2n* | ACOL |  | 46 | 6 | 9 | 0.704 | 0.00201 | -0.77528 | 0.33477 | -0.01513 |
|  | AGAM |  | 34 | 2 | 3 | 0.658 | 0.00168 | 1.26352 | 0.79033 | 1.07038 |
|  | AFUN |  | 80 | 5 | 6 | 0.668 | 0.00197 | -0.15261 | 1.05574 | 0.78249 |
|  | All |  | 160 | 7 | 11 | 0.673 | 0.00192 | -0.57314 | 1.15765 | 0.66449 |
| Abdomen (oocyst) +<br>Head/Thorax<br>(sporozoite) | ACOL |  | 104 | 8 | 13 | 0.683 | 0.00203 | -1.08841 | -0.95047 | -1.18445 |
|  | AGAM |  | 66 | 3 | 4 | 0.621 | 0.00161 | 0.42848 | 0.86413 | 0.85268 |
|  | AFUN |  | 176 | 6 | 8 | 0.646 | 0.00192 | -0.53471 | 1.15025 | 0.67236 |
|  | All |  | 346 | 9 | 17 | 0.652 | 0.00189 | -0.92186 | 1.28984 | 0.56723 |
| Whole mosquito;<br>2n* (mixed oocyst +<br>sporozoite stages) | AFUN |  | 40 | 2 | 3 | 0.610 | 0.00149 | 0.97432 | 0.77124 | 0.96165 |

**Fig. 3.**
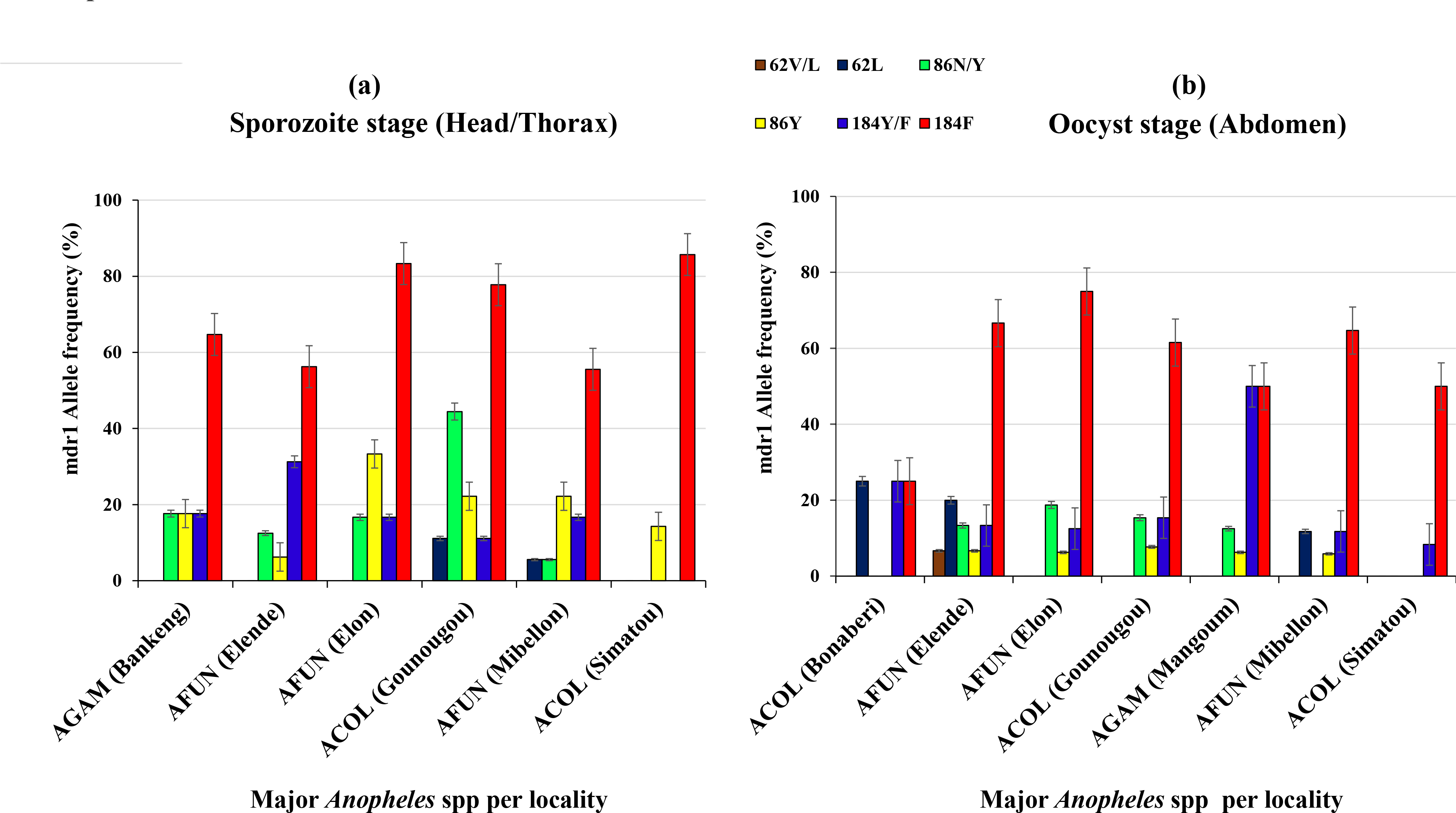

### 3.4. Genetic variability of the *Pfk13* propeller gene in major *Anopheles* vectors across Cameroon

Analysis of the 789-bp fragment of the *k13* propeller domains from 88 *P. falciparum* oocyst infected *Anopheles* abdomen sequences revealed the existence of ten distinct haplotypes (Table 3a). Generally, eight (8) polymorphic sites were detected with a haplotype diversity of 0.163. The dominant wild haplotype (H1) scored a high frequency of 91.5% (161/176) (Fig. 4a & 4b) while the remaining haplotypes constituting the mutant sequences recorded a low frequency including the H2 (0.57%, 1/176), H3 (1.14%, 2/176), H6 (0.57%, 1/176), H7 (0.57%, 1/176), H8 (2.28%, 4/176), H9 (0.57%, 1/176) and H10 (1.14%, 2/176) in *An. funestus*. The H4 (0.57%, 1/176) and H5 (1.14%, 2/176) haplotypes were documented in *An. gambiae* (Fig. 4b). Sequences congregate according to the presence of mutation, with the sequences containing mutations distancing away from sequences of H1. Deviation from the neutrality test of Tajima’s D had a significant negative value (D= -1.91573*) indicating an excess of rare alleles in the population (Table 3a). A maximum likelihood (ML) tree of the sequences analyzed confirms the low diversity with three major clusters (Fig. 4a). The wild type sequences dominated as the most representative group followed by two mutant allele clusters in *An. funestus* (Fig. 4a). The remaining discrete cluster represents the mutant *P. falciparum* sequences from *An. gambiae* and *An. funestus.* Moreover, the haplotype network tree analysis revealed that haplotypes H2 to H10 are separated by a single mutation step from the ancestral haplotype H1 (Fig. 4b).

**Fig. 4.**
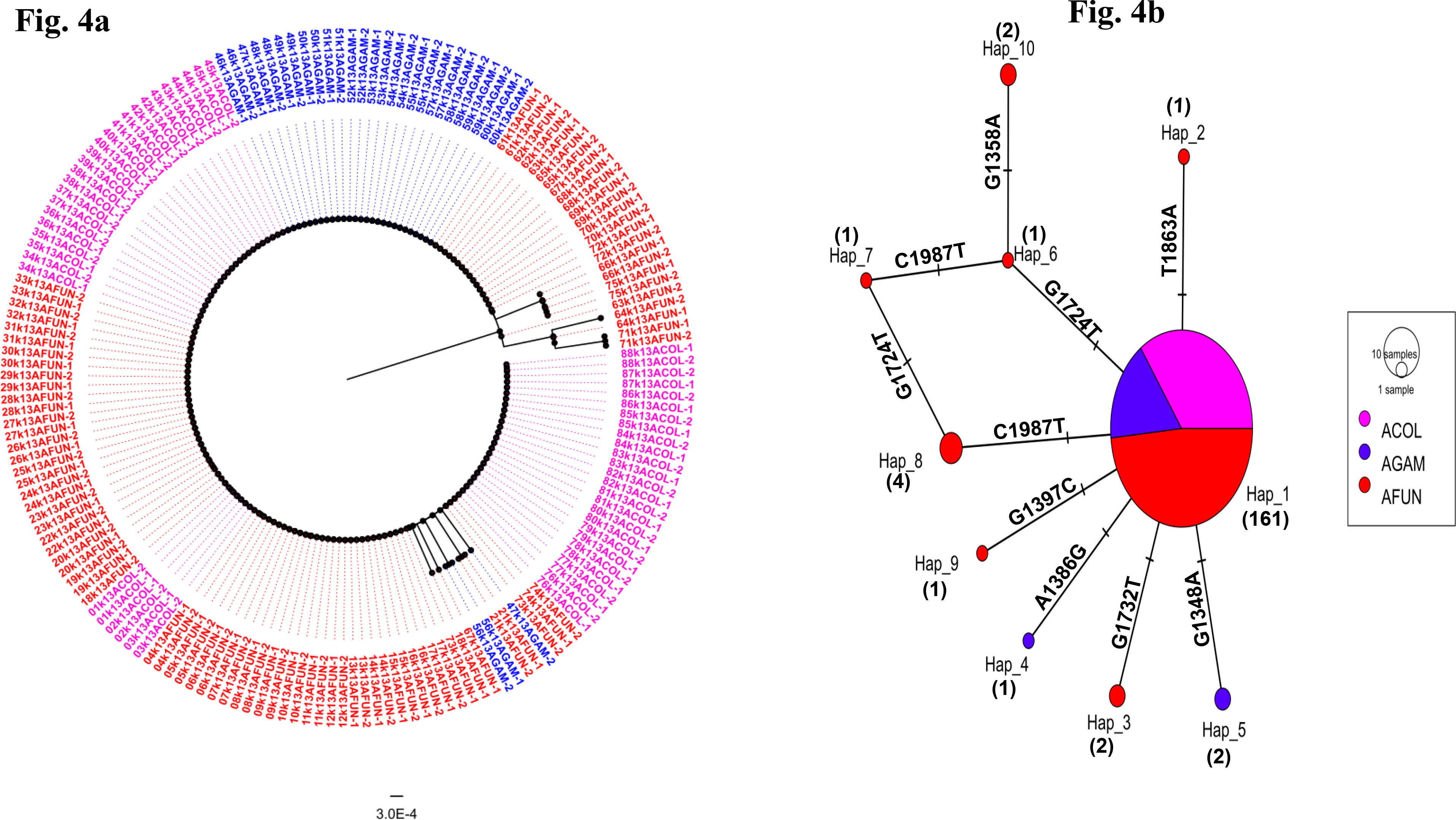

Polymorphism analyses of *P. falciparum* sporozoites in the H/T of infected *Anopheles* mosquitoes across Cameroon revealed a reduced haplotype diversity of 0.113 and five (5) polymorphic sites (Table 3a). Six (6) haplotypes were identified with the superior haplotype, H1 (94.2%, 130/138) representing the wild type population backbone. The remaining haplotypes representing the mutant sequences recorded a low frequency including: H2 (1.45%, 2/138), H3 (1.45%, 2/138), H5 (1.45%, 2/138) and H6 (0.72%, 1/138) in *An. funestus* from Elende; H4 (0.72%, 1/138) in *An. coluzzii* (Fig. 5a & 5b). The R575I mutation was absent in the H/T sporozoite sequences. Also, a reduction in the number of *k13* haplotypes between the oocyst (abdomen; n = 10) and sporozoite (H/T; n = 6) (2 = 3.01; *p* = 0.02) was observed. Phylogenetic χ analysis confirmed three distinct haplo-groups with the major haplotype being the wild-type sequences and the other bearing the five minor mutant sequences (Fig. 6a). Furthermore, the neutrality population inference statistic of Tajima D (D=-1.76645), FuLiD (-1.19624) and FuLiF (-1.63960) tests were all negative possibly indicating presence of rare alleles driven by a strong selection pressure (Table 3a). Similarly, the separation of H2 to H6 minor haplotypes from the parental major haplotype by one mutational line (Fig. 5b) highlights the independent emergence of these alleles. On the other hand, maximum likelihood and haplotype analysis of 28 *P. falciparum* isolates in *An. funestus* whole mosquito sequences from Obout showed the presence of only one polymorphic site and a single haplotype comprising of the wild type sequences. Tajima D and FuLiD variables were all zero (Table 3a).

**Fig. 5.**
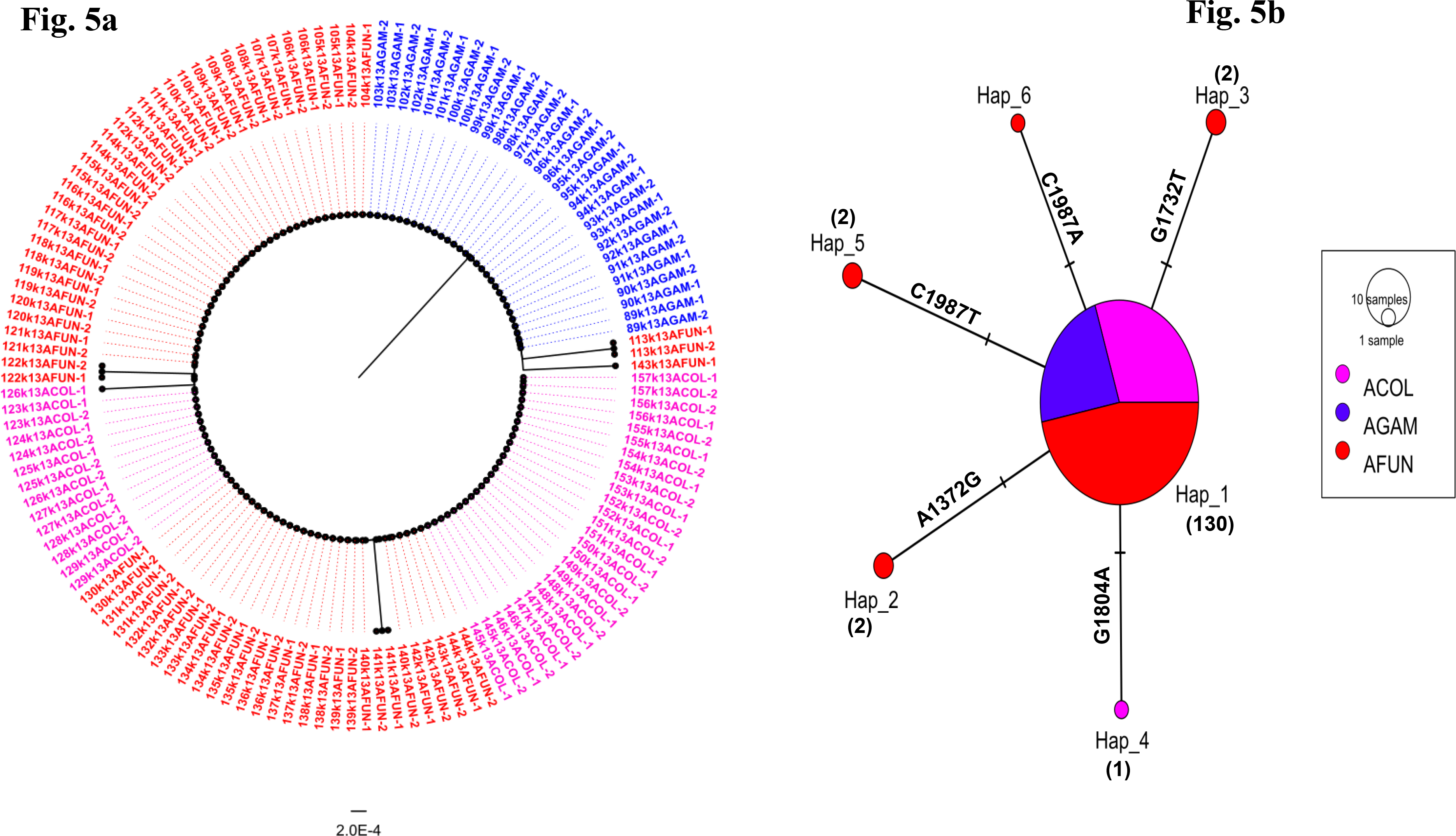

**Fig. 6.**
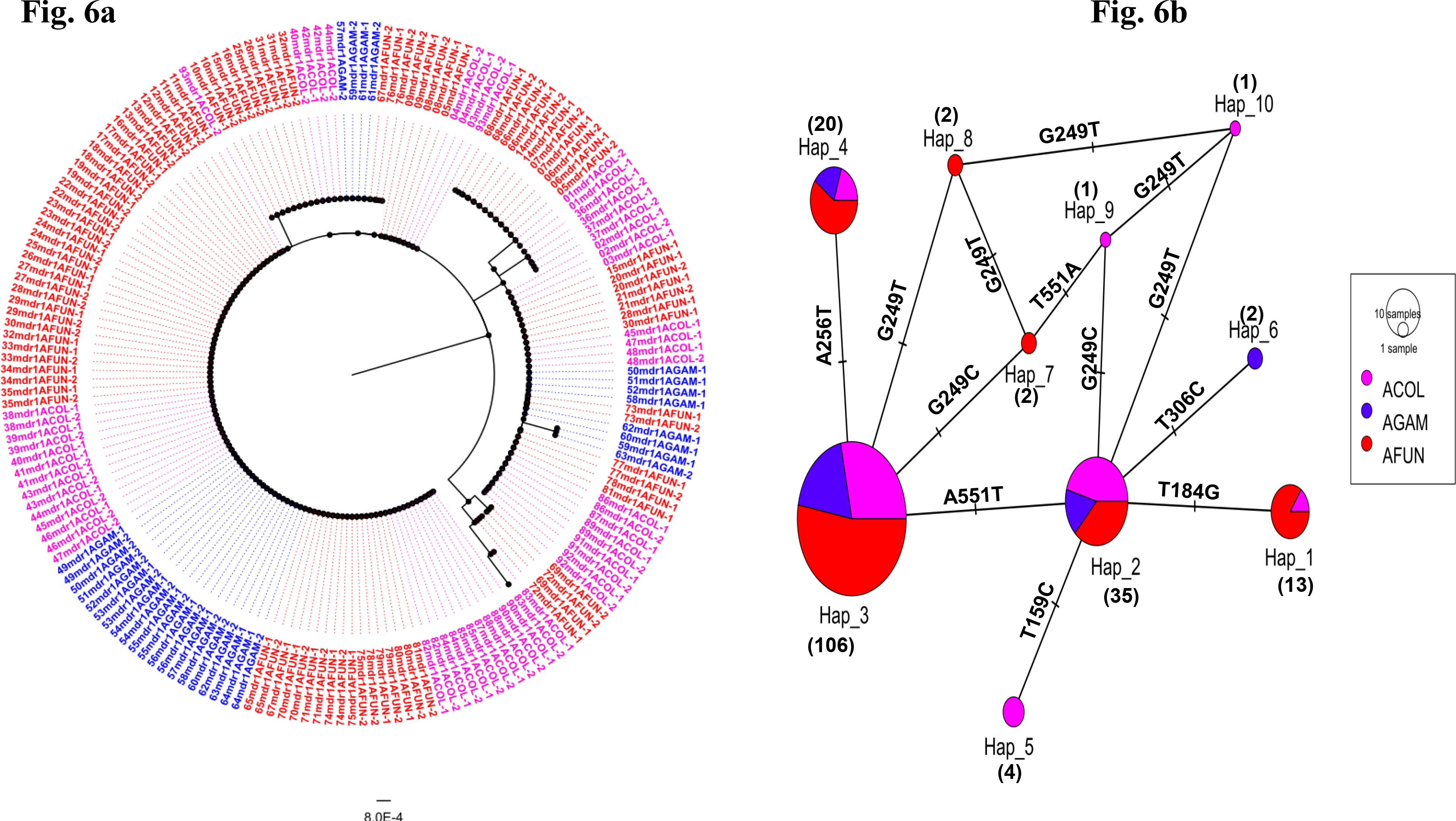

### 3.5. Polymorphism analysis of the *Pfmdr1* gene fragments in the major *Anopheles* vectors across Cameroon

Six polymorphic sites (6) and ten haplotypes were detected across seven localities. The haplotype diversity was 0.626 (Table 3b). The predominant haplotype, H3, comprised exclusively of populations harboring the 184F resistant allele backbone occurring at a frequency of 56.9% (106); followed by the H2 haplotype representing only the wild type allele at a proportion of 18.8% (35) (Fig. 6b). Haplotypes, H1 (6.9%, 13) comprises only the 62L allele while H4 (10.8%, 20) is a mix of the 86Y and the 184F resistant variant backbones. Co-existing haplotypes including the H5, H6, H7, H8, H9 and H10 occurred at a minor frequency of 2.2% (4), 1.1% (2), 1.1% (2), 1.1% (2), 0.54% (1) and 0.54% (1) respectively (Fig. 6b). Proximity to the neutrality statistic of Tajima’s D had a negative score (D= -0.59191) indicating both population expansion and emergence of rare alleles owing to strong ACT pressure. A phylogenetic tree plot of the sequences circulating in the abdomen reveals six major clusters with the key 184F, 86Y and 62L backbones each forming a group (Fig. 6a). The haplotype network tree shows that the mutation emerged as a result of parasite gene flow events facilitated by human and vector mobilities. This is evident based on the fact that shared haplotypes were observed in all the three major *Anopheles* vectors (Fig. 6a).

Polymorphism analyses of unphased *P. falciparum* sporozoites in the H/T of infected *Anopheles* mosquitoes in six localities across Cameroon revealed a haplotype diversity of 0.673 with eight polymorphic sites (Table 3b). A total of eleven (11) haplotypes were identified with the major haplotype H1 (48.8%, 78/160) solely harboring the 184F resistant backbone (Fig. 7b). H2 (21.9%, 35) was shared between 184F and 86Y alleles and H3 (21.3%, 34) was a mixture of wild type and the mutant haplotypes (86N and 184F). Haplotypes, H5, H6 and H9 harbored the 62L allele (2.5%, 4) while H4, H7, H8, H10 and H11 occurred at minor frequencies of 1.3% (2), 0.6% (1), 1.3% (2), 1.3% (2) and 1.3% (2) respectively (Fig. 7b). Phylogenetic analysis confirms four distinct haplotype groups with the main haplotype being the 184F resistant assemblage alongside the one sub-major 86Y resistant parasites backbone (Fig. 7a). Furthermore, the neutrality population inference statistic of Tajima D (D=-0.57314) was negative alongside a positive FuLiD (1.15765) and FuLiF (0.66449) tests (Table 3b). As previously indicated, the negative Tajima D implies a significant expansion of the mutant parasite population driven by a strong selection pressure. Similar results were observed for the whole *P. falciparum* infected *An. funestus* population (Table 3b) with three major haplotypes observed (Fig. 8a & 8b). Further analysis combining both the oocyst (abdomen) and sporozoite (H/T) population reveals 17 circulating haplotypes with 07 distinct phylogenetic groups (Table 3b).

**Fig. 7.**
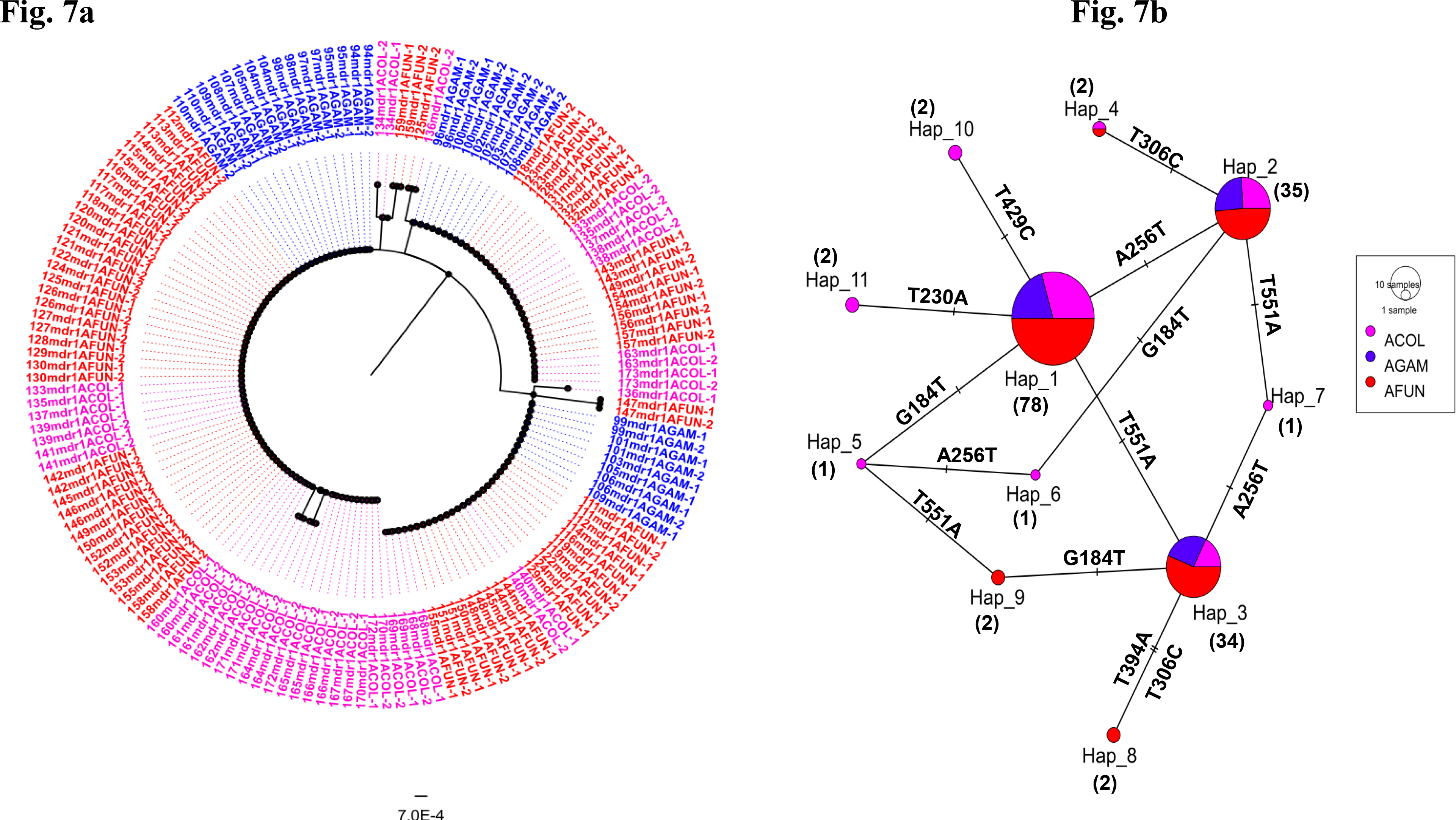

**Fig. 8.**
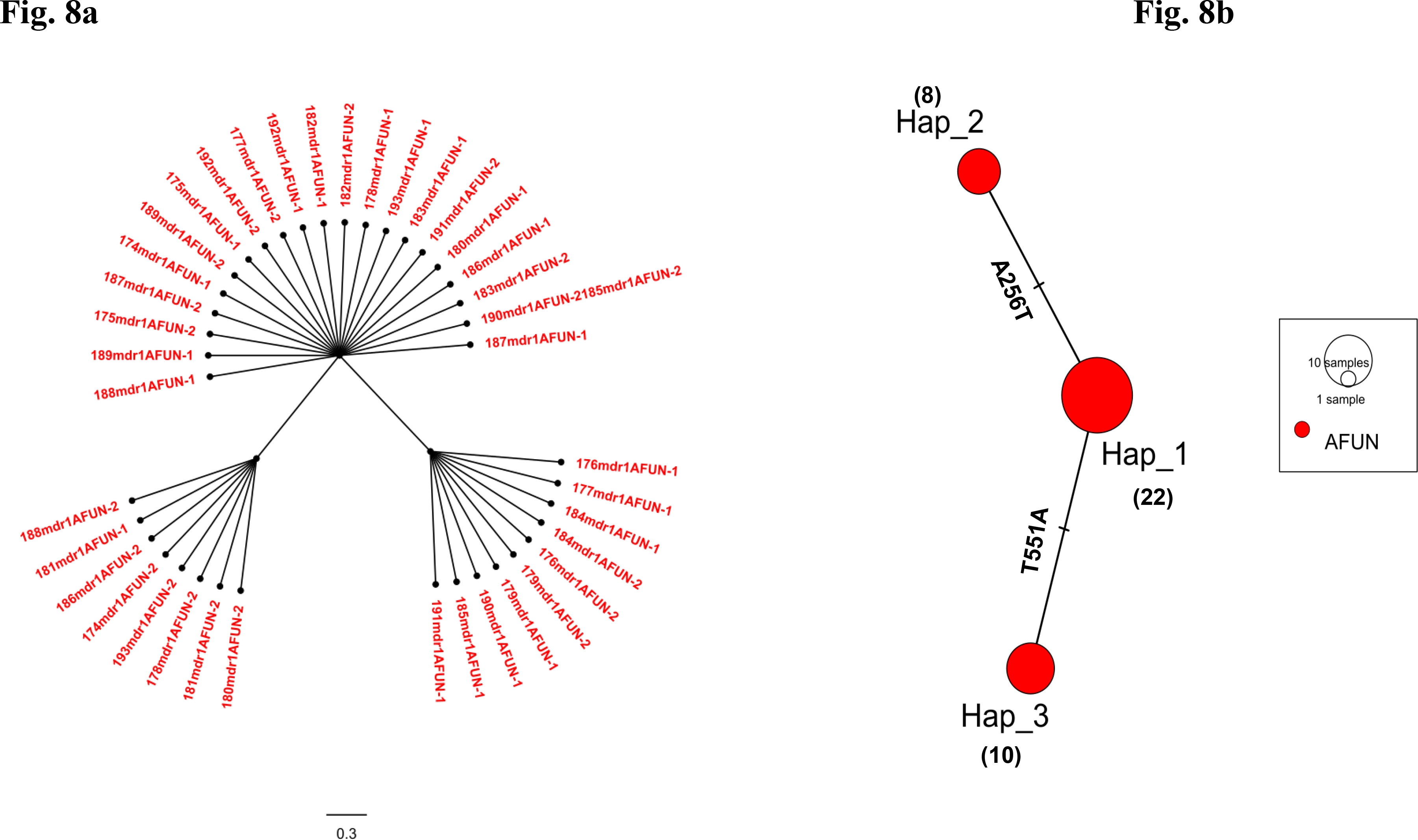

## 4. Discussion

In the pursuit to control malaria, understanding the processes governing transmission of parasites between *Anopheles* vector and humans is fundamental [50]. This is particularly relevant in the Cameroon context where heterogeneous malaria transmission across diverse bioecological landscapes [3] is driven by many factors including the *Plasmodium* infection rate in genetically diverse *Anopheles* vectors. To this effect, anti-malaria drugs are heavily deployed to tackle the parasite population in an effort to minimize transmission. Such massive intervention of anti-malaria drugs is the major driving selection force of parasite evolutionary adaptation particularly for *P. falciparum* [51]. This ultimately leads to the emergence of drug resistance alleles which are transmitted by *Anopheles* vectors to humans during an infectious blood meal. Currently, surveillance for *P. falciparum* markers mediating resistance to artemisinin and partner drugs is mostly based on parasite genotyping from positive human blood samples by microscopy [22]. However, naturally infected mosquitoes are rarely utilized despite being the definitive host where sexual recombination occurs, producing a mixture of phenotypes. Here, genomic phenotyping of *P. falciparum* DNA from field-caught *Anopheles* mosquitoes across Cameroon was performed to investigate the possible emergence of mutations involved in artemisinin resistance and to determine the frequency of the *mdr1* alleles implicated in partner drug tolerance. This study provides data on the *Plasmodium* infection rate and polymorphism profile of molecular markers mediating drug resistance in *P. falciparum* infected *Anopheles* mosquitoes driving malaria transmission across Cameroon.

### 4.1. High *Plasmodium* infection rates of major *Anopheles* vectors across Cameroon suggest a heterogenous malaria transmission across diverse ecological biotopes

Despite the widespread distribution and overall impact of long-lasting insecticide treated nets, there still exists a significant variation in *Plasmodium* infection prevalence in the major malaria vectors across Cameroon [3, 32]. The infection rate which is a key component of the entomological inoculation index determines the local patterns of malaria transmission [3]. The heterogeneity in the observed *Plasmodium* infection rates per each eco-geographical landscape translates to the vectorial competence of the dominant vectors. Villages located within the equatorial facets (Bankeng, Elende, Mangoum and Obout), sudano-guinean belt (Mibellon and Gounougou) and Sahelian zone (Simatou) where *An. funestus*, *An. gambiae* and *An. coluzzii* are generally the principal vectors recorded high sporozoite infection rates. This could be as a result of high vector densities [5], high degree of pyrethroid resistance that encourages the tremendous anthropophilic blood feeding tendency of the vectors and their ability to survive in nature [32, 52]. These factors favor the vectors’ ability to potentially blood feed on the humans harboring high *Plasmodium* parasite loads thereby piloting the hyper-endemic malaria transmission pattern observed in these localities. However, a low sporozoite infection rate was observed in *An. coluzzii* population from Bonaberi where malaria transmission is hypo-endemic. This disparity could be attributed to a low parasite prevalence in the resident human population [53] and the transmission capacity of the local vector [54]. Moreover, the extensive circulation and use of drugs in this urban setting may contribute to shrinking the parasite population pool in humans [55]. In particular, the high sporozoite infection rate scores of *An. funestus* provides further evidence on the major role played by this vector species in determining malaria transmission in Cameroon. This observed infection rate is similar to previous reports for this species in Cameroon [32] and across Africa precisely in Benin (18%) [56] and Ghana (12.5%) [57]. This repeatedly high score of infection corresponds with a high vectorial capacity of this *Anopheles* species across Africa and poses a serious concern on the efficacy of interventions targeted at controlling malaria in high transmission areas.

### 4.2. The impact of mosquito intrinsic defense system and midgut barriers in reducing the force of *Plasmodium* infection and malaria transmission

The marked reduction in the sporozoite infection rate relative to the oocyst infection rate notably for *An. funestus* population from Mibellon, Elon and Elende and *An. coluzzii* population from Bonaberi indicates the contribution of mosquito immune system checkpoints [58] in limiting both *P. falciparum* and *P. malariae* sporogonic development. Thus, exploiting mosquito immunity for possible design of transmission blocking interventions will complement efforts towards malaria elimination in Africa.

### 4.3. Absence of artemisinin resistance markers in *P. falciparum* populations suggest the continued efficacy of artemisinin derivatives in Cameroon

The absence of the WHO validated *Pf*k13 mutations (F446I, Y493H, R539T, I543T, P553L, R561H, P574L, C580Y, A675V, C469Y) [1, 22, 23, 59] possibly reflects the continued efficacy of artemisinin derivatives in the treatment of malaria in Cameroon. Majority of SNPs in the *k13* protein of *P. falciparum* parasite population from African descent revolves particularly within the 400-600 amino acid coding region of the β ropeller domain [25, 59], indicating that this region is particularly under strong selection owing to among others, intense ACT use over the years [21]. This points attention to the R575I polymorphism detected at a low frequency in the oocyst (abdomen) of *An. funestus* mosquitoes from Mibellon. This mutation, previously observed in Rwanda at a much lower frequency has not yet been characterized [22], although it was observed not to be associated with artemisinin resistance in Rwanda parasite isolates [22]. However, this R575I SNP is located five amino acids upstream of the C580Y and is also adjacent to the R56IH mutation which are the key variants mediating Art-R in South-East Asia and Rwanda respectively. Secondly, like the R561H and C580Y SNPs [22, 59], the R575I mutation is interesting because Isoleucine, a neutral non-polar amino acid replaces Arginine, a basic polar amino acid. This substitutional change may affect the tertiary structure and thus the function of the propeller [60]. Nonetheless, the absence of this mutation in the sporozoite (H/T) of infected *An. funestus* mosquitoes may imply that mutation-induced fitness could be playing a vital role in limiting the transmission of mutant parasite phenotypes [61]. *P. falciparum* parasites harboring deleterious artemisinin resistant alleles could be less fit for survival compared to wild types and the probability of their transmission is reduced. This is particularly relevant for rare *k13* mutations that may impose a huge survival cost for the parasite especially in high malaria transmission settings where competition with multiple parasite strains is common [51]. This could explain the reason why the R575I mutation was present in the oocyst but absent in the sporozoite stage. This study highlights the viewpoint that regular monitoring for *de novo* emergence of potentially novel local variants and surveillance for the possible introduction of the resistance phenotype is necessary for tracking the emergence of artemisinin resistance and mapping resistance hotspots in Cameroon.

Furthermore, the A578S variant, commonly observed in different *P. falciparum k13* backgrounds across Africa has already been demonstrated not to be involved in artemisinin resistance [59, 62]. This polymorphism was detected at a low frequency in the abdomen and H/T of two sperate *An. funestus* mosquito from Elon. A recent study evaluating the clinical efficacy of dihydroartemisinin/piperaquine in *P. falciparum* malaria subjects in Yaoundé (situated 50 Km from Elon), Centre Cameroon revealed a 2% (3/150) prevalence of the A578S mutation [63], further confirming the data obtained from this study. However, comparing previous studies in Cameroon that utilized *P. falciparum* infected blood samples from human subjects [64, 65], none of the *k13* polymorphisms detected in this study was similar. This could be attributed to many factors including, the wide variability of the polymorphism profile in the *k13* region, differences in parasite genotype complexity, sampling sites, period of sampling, local selection pressure, intensity of malaria transmission and level of immunity [51]. Moreover, the fact that these three studies utilized *P. falciparum* human blood samples as opposed to infected *Anopheles* mosquitoes for molecular genotyping could reflect dissimilarities in both the human (intermediate) and *Anopheles* (definitive) host systems wherein parasites harboring lethal mutations may be eliminated during the sexual development in the mosquito. Also, the presence of the A578S polymorphism in both oocyst and sporozoite stages suggest a neutral effect and no fitness cost of this mutation associated with parasite transmission [61]. The significant negative D value suggests a possible recent *de novo* expansion of singleton polymorphisms in this gene across the parasite populations. No evidence of selection acting on the domain implies that ACTs pressure is minimally impacting on *k13*PD diversity in Cameroon. Generally, the *k13*PD locus exhibits a remarkable genomic sequence conservation across *Plasmodium* species [66] and the fact that no fixed non-synonymous mutation was found in the domain indicates that this gene evolved under strong purifying selection; and that the rare variants observed in this study arose recently. Indeed, a large number of rare variants is characteristic of the genomes of African but not Asian *P. falciparum* parasites [66].

### 4.4. Directional selection of Y184F in combination with the N86Y allele alongside emerging novel variants may be compromising parasite susceptibility to ACTs

Mutations in the *Pfmdr1* gene is associated with decrease sensitivity to multiple antimalarial drugs including, amodiaquine, lumefantrine, halofantrine, mefloquine, chloroquine and artemether [26, 67, 68]. The high frequency of the ^86^N^184^F alleles implies that *Pfmdr1* parasite mutants are selected and sustained in the population. This could be linked to the continuous deployment of first-line interventions; Artemether-Lumefantrine (AL) and Amodiaquine-Artesunate (ASAQ) for the treatment of uncomplicated malaria [3]. These observations correlate with similar findings in Cameroon where blood samples from human cohorts were used for resistance genotyping [69] and elsewhere in Africa where infected mosquitoes were the targeted group [27, 70]. A greater fraction of the parasites harbored the 184F allele at both oocyst (60.2%) and sporozoite (67.5%) stages. However, a disparity was observed for the 86Y resistant variant where a significant proportion (χ^2^ = 3.39, *P* 0.05) of this allele was harbored by *P. falciparum* sporozoite population (17.5%) as compared to the oocyst population (5.4%). These differences reflect the opposing effects of AL and ASAQ. Indeed, studies have demonstrated that AL selects for both the N86-wild type and 184F-mutant haplotypes (^86^N^184^F) whereas ASAQ is associated with selection of the mutant 86Y and wild Y184 haplotypes (^86^Y^184^Y) [71]. The predominance of the AL selected NF haplotype suggests its continuous selection over time; implying that these mutant parasites have better adapted and are more fit for transmission, and may constitute the dominant population involved in gametocytogenesis.

Contrary to *k13* mutant phenotypes, parasites harboring the *mdr1* allele, Y18F may have a significant advantage due to the long-standing existence of resistance to the partner drug, further facilitated by decades of continuous drug use [67, 71]. This creates an even stronger sustained selection pressure over time, consequently favoring transmission of these mutant parasite phenotypes. The detection of the V62L variant points to the view that the *mdr1* gene backbone could be evolving in some localities via the emergence of novel alleles as ACTs are continuously implemented. Nevertheless, the reduction in the frequency of the V62L novel allele from the oocyst to sporozoite stage notably in Mibellon and Elende *P. falciparum* infected *An. funestus* populations may reflect a possible fitness cost to enhanced clearance by mosquito immune mechanisms.

### 4.5. Mosquito immune selection may be suppressing the spread of artemisinin drug resistance parasites while facilitating the transmission of AL selected parasite lineages

African malaria vectors particularly *An. gambiae* and *An. funestus* has been shown to mount high levels of innate immune response against the *Plasmodium* parasite especially after sexual recombination in the midgut [72]. Such intensity in the strength of immune selection may limit the survival of lethal *P. falciparum* drug resistant variant phenotypes while concomitantly bringing about a bottleneck reduction in oocyst loads [73]. In particular, genetic recombination can negatively impact the spread of artemisinin resistance alleles by breaking down resistance haplotypes [28]. This therefore implies that mosquito immune selection may be delaying the development and spread of potentially dangerous drug resistance alleles in high transmission settings by favoring sensitive lineages to remain in the hemocoel circulation, particularly for *k13* polymorphisms involved in artemisinin resistance. In addition, the interspecific competition within genetically diverse parasite phenotypes may lead to the suppression of resistant lineages within *Anopheles* hosts [28].

However, as the occurrence of sexual recombination in the mosquito midgut produces a mixture of phenotypes; this may positively influence the spread of partner drug resistant alleles through transmission dependent mosquito selection of a particular haplotype known to potentiate parasite tolerance (e.g., ^86^N^184^F for AL). Such traditionally resistant genetic variants may confer better adaptative potentials for parasite transmission. In addition, once resistance is common on many *P. falciparum* genetic backgrounds (notably for *mdr1*), mosquito immune selection can steadily maintain a particular resistance allele of the parasite to the existing drug (e.g., lumefantrine or amodiaquine) in the population. This eventually favors the multiplication and transmission of the resistant parasite phenotype. This could account for the dominance of the 184F resistant allele present in *P. falciparum* infected mosquitoes and reveals that this allele is under transmission-driven positive selection. This suggests that mosquito immune selection may be shaping the local pattern of drug resistance evolution and spread once resistance is widespread [74]. This agrees with a previous study demonstrating that mosquitoes play a significant role in determining the frequency of drug resistant *P. falciparum* population in natural [28, 29, 70] and experimental [75] settings.

Hence, in high malaria transmission settings, wild mosquitoes could be playing a key role in delaying the emergence and spread of artemisinin drug resistant parasites. Also, this study points to the view that interventions focusing on reducing mosquito abundance may indirectly influence the spread of drug resistant parasites by decreasing the *Anopheles* population that select on drug resistance polymorphisms [28]. Additional investigation is required to assess the impact of mosquito immunity on the development and transmission of drug resistant parasites through experimental infection assays.

Although drug resistance xeno-surveillance of *P. falciparum* from the lens of the mosquito is a promising approach, there exist a few drawbacks. Firstly, detection of emerging selected *de novo* mutants linked particularly to artemisinin resistance could be more sensitive in the human host, contrary to the mosquito life cycle, where sexual recombination driven-negative selection against such mutants could occur [76]. In spite of this, allele frequency estimate of resistance marker surveillance in the mosquito vector is reliably a more appropriate measure of resistance epidemiology since anti-malaria drug resistance presents a higher chance to be transmitted than acquired [27]. Furthermore, these infected mosquitoes were mostly collected in high transmission settings, implying that xeno-monitoring approach may be less economical in a low malaria transmission area (e.g., Bonaberi) where screening of a considerable large number of mosquitoes has to be undertaken to detect both low-density infections and polymorphism counts. Notwithstanding, this is a common problem for resistance detection in low transmission settings, regardless of utilizing human or mosquito samples.

## 5. Conclusion

Xeno-monitoring provides an alternative practical approach to the surveillance of *Plasmodium* parasite populations in *Anopheles* vectors, especially in high transmission settings. The high *P. falciparum and P. malariae* infection rates in *Anopheles* mosquitoes coupled with the dominance of the ^86^N^184^F haplotype suggest an increase tolerance of *P. falciparum* to ACTs particularly Artemether-Lumefantrine and highlights the challenges of malaria control in Cameroon.

## Abbreviations

ACTs: Artemisinin Combination Therapy; AL: Artemether-Lumefantrine; ASAQ: Amodiaquine-Artesunate; ITNs: Insecticide-Treated Nets; NMCP: National Malaria Control Program; Art-R: Artemisinin Resistance; ELISA: Enzyme-Linked Immunosorbent Assay; HLC: Human Landing Catch; PSC: Pyrethrum Spray Catch; *k13*PD: Kelch 13 propeller domain; *mdr1*: multi-drug resistance-1; PIR: *Plasmodium* infection rate; P.OVM: *Plasmodium ovale*, *vivax* and *malariae*; Spz: Sporozoite; WHO: World Health Organization;

## Ethical approval

This study received institutional approval (N^0^ 2020/05/1234/CE/CNERSH/SP) for HLC sampling from the Cameroon National Committee on Research Ethics for Human Health.

## Data Availability Statement

All the data from this study is present in the manuscript and supplementary file (Appendix A). All *Pfk13* and *mdr1* sequences in this study were deposited in the GenBank database (accession numbers - *Pfk13*: OM023056 - OM023397 and *Pfmdr1*: OM023398 - OM023783).

## Competing interest

Authors declare no conflict of interest.

## Funding statement

This work was funded by the Wellcome Trust Senior Research Fellowship in Biomedical Sciences (Grant No. 217188/Z/19/Z) awarded to CSW. The funders had no role in the design, data collection, analysis, interpretation of the results, preparation of manuscript or decision to publish.

## Author contributions

C.S.W. conceived and designed the study. F.N.N., L.M.J.M., and M.T. (Magellan Tchouakui) collected some of the mosquito samples on the field. F.N.N. performed mosquito DNA extraction with assistance from L.M.J.M and M.T. (Magellan Tchouakui). F.N.N. performed the TaqMan experiments and drug resistance genotyping. M.J.W., B.M., M.T. (Micareme Tchoupo) provided previous mosquito collections. F.N.N., M.J.W., B.M. and M.T. (Micareme Tchoupo) handled the sequencing preparation at CRID. F.N.N., L.M.J.M., and N.N.D. performed sequence analysis. C.S.W. provided the resources and validated the data. ., S.W., and C.N. supervised the study. F.N.N. and C.S.W. wrote the manuscript with contribution from all the authors. All authors read and approved the final version of the manuscript.

## Acknowledgments

The authors are grateful to the research team members at CRID particularly Doumani Djonabaye, Dr Delia Djuicy and Yvan Fotso for their scientific and graphical inputs and to Mr. Achille Binyang for providing the Bankeng mosquito samples. We also thank the village leaders and community members of the various localities for providing access to their houses to be used for mosquito sampling.

## Notes

### Competing Interest Statement

The authors have declared no competing interest.

